## Supplementary material for "Primosomal protein PriC rescues replication initiation stress by bypassing the DnaA-DnaB interaction step for DnaB helicase loading at *oriC*": Figure supplements

### Figure 2-figure supplement 2

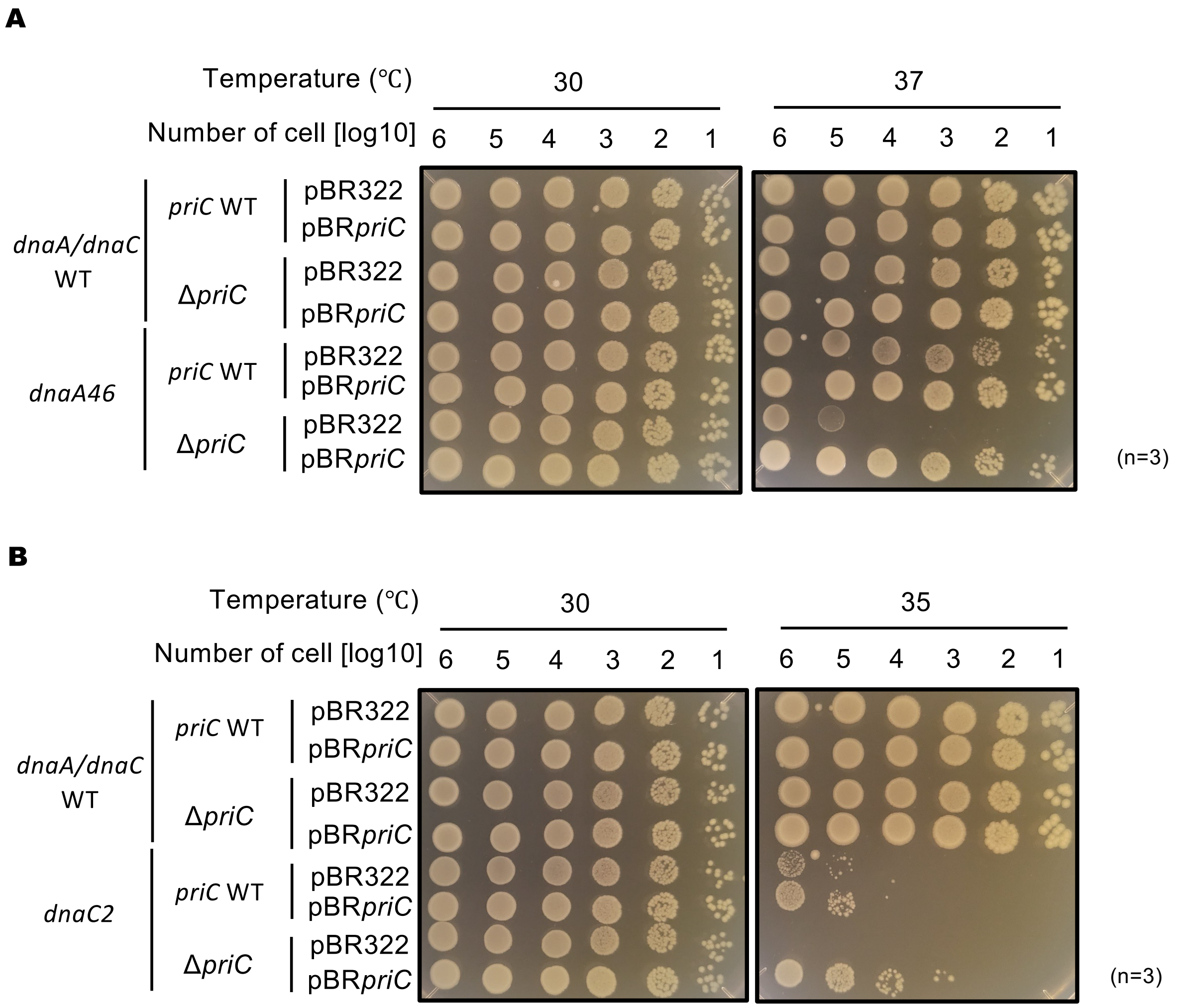

### Figure 2-figure supplement 2

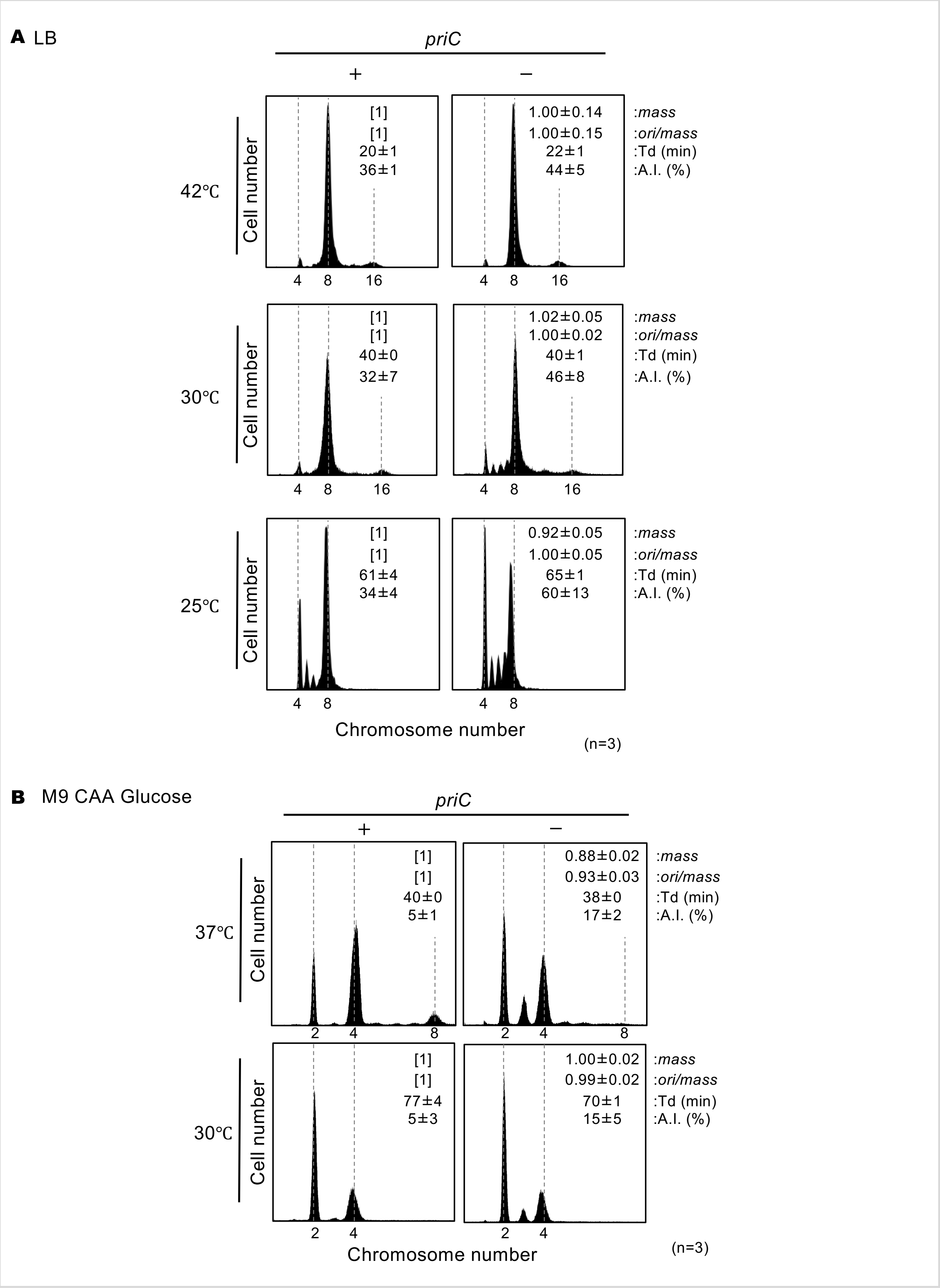

### Figure 3-figure supplement 1

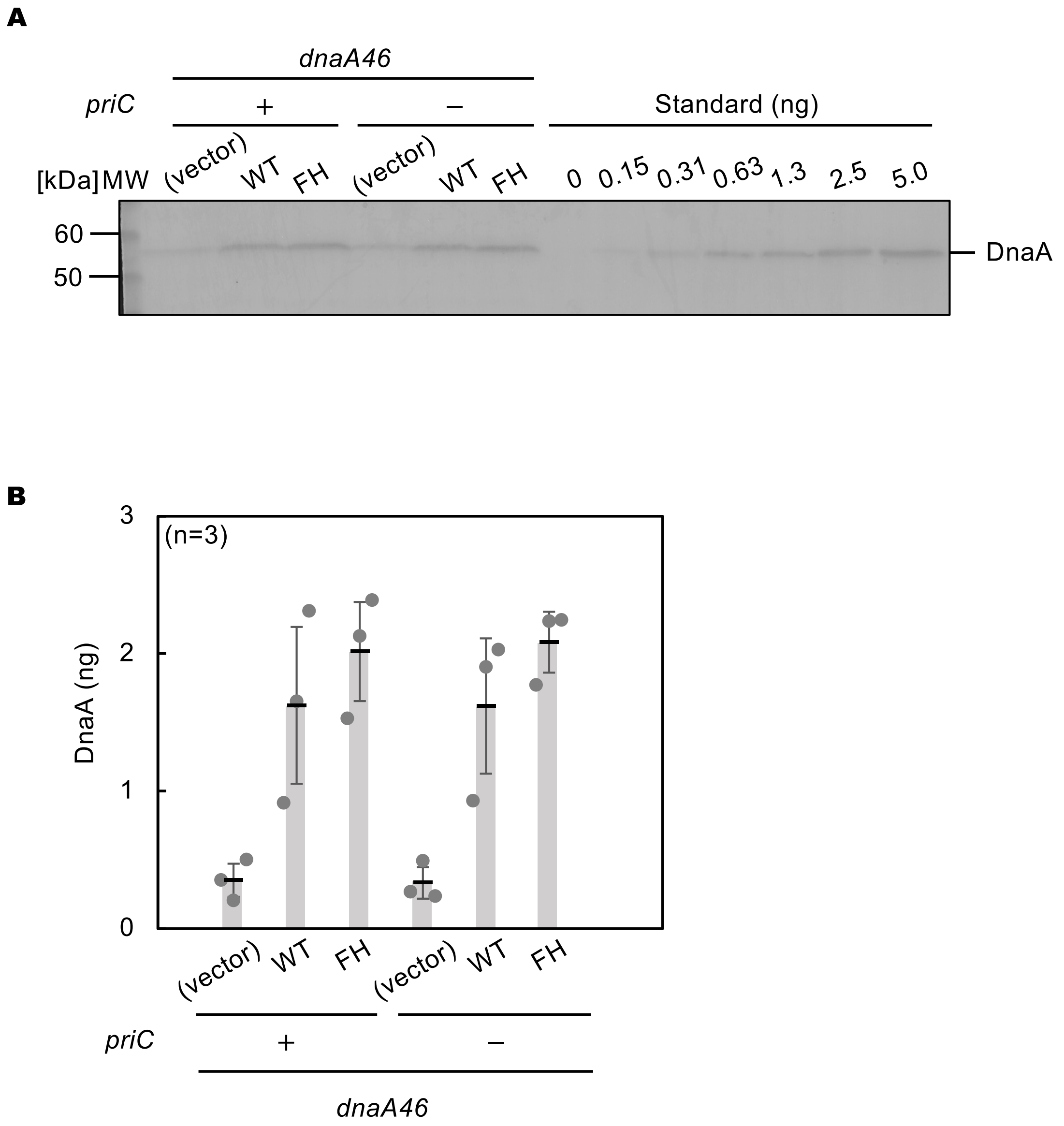

### Figure 4-figure supplement 1

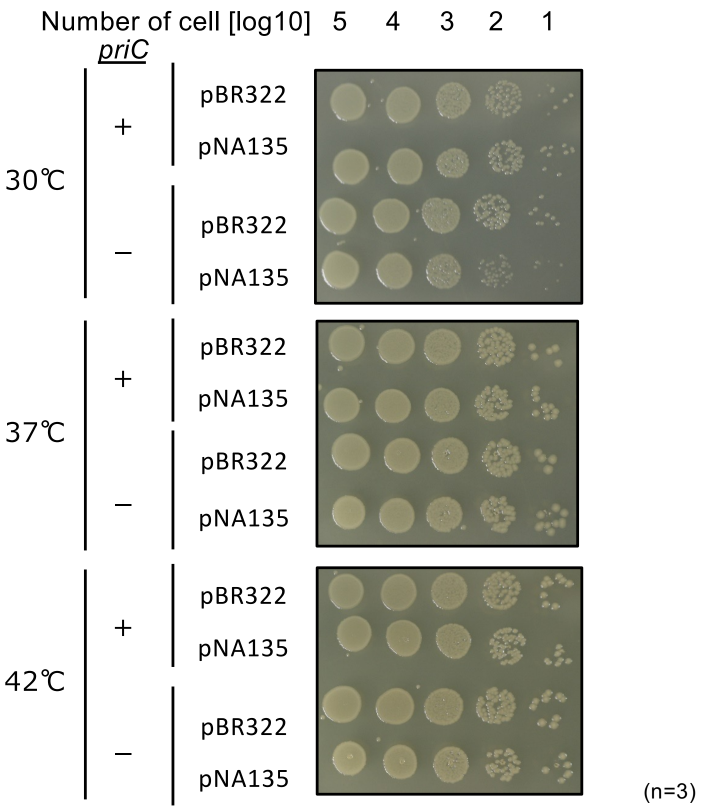

### Figure 7-figure supplement 1

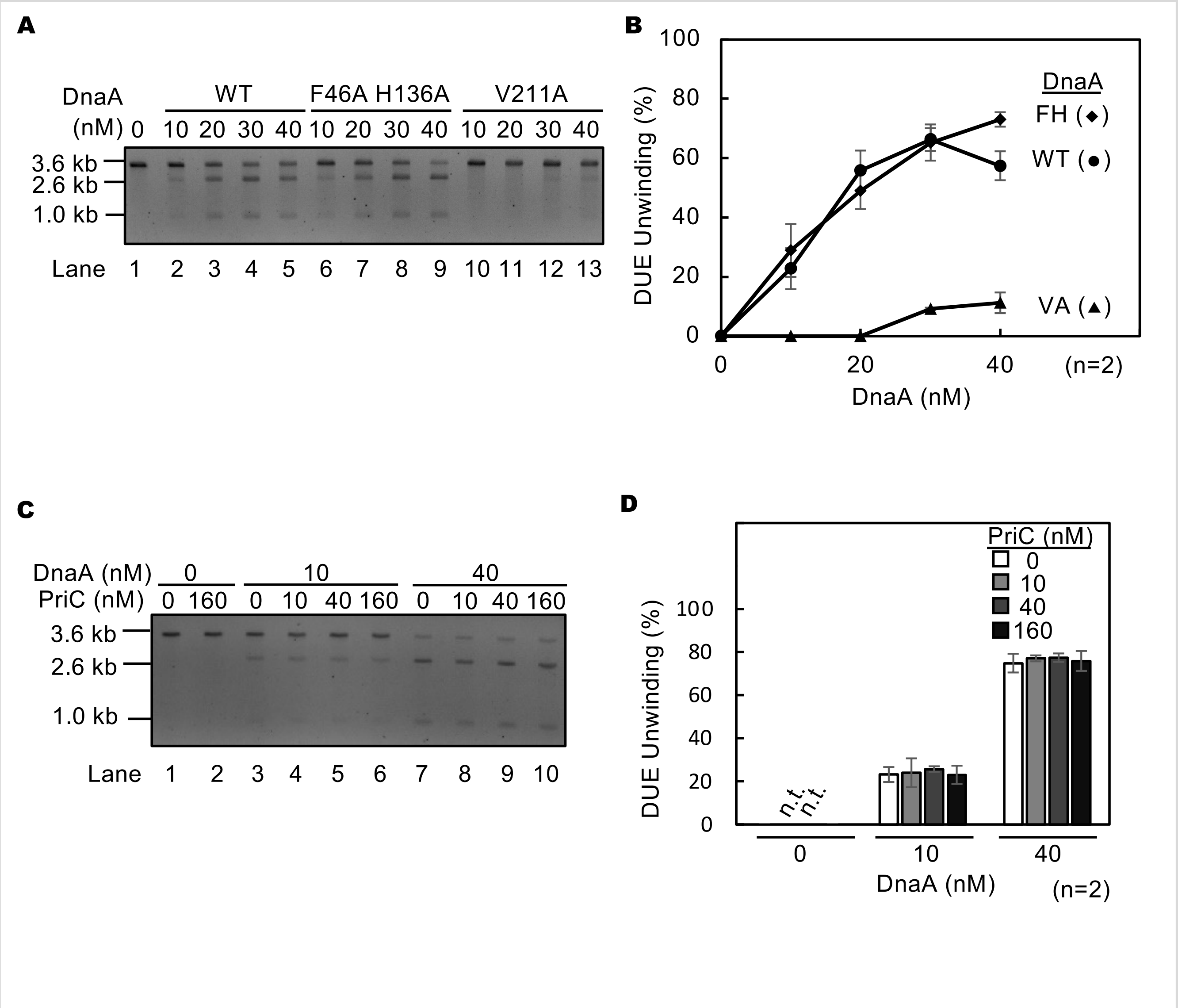

### Figure 8-figure supplement 1

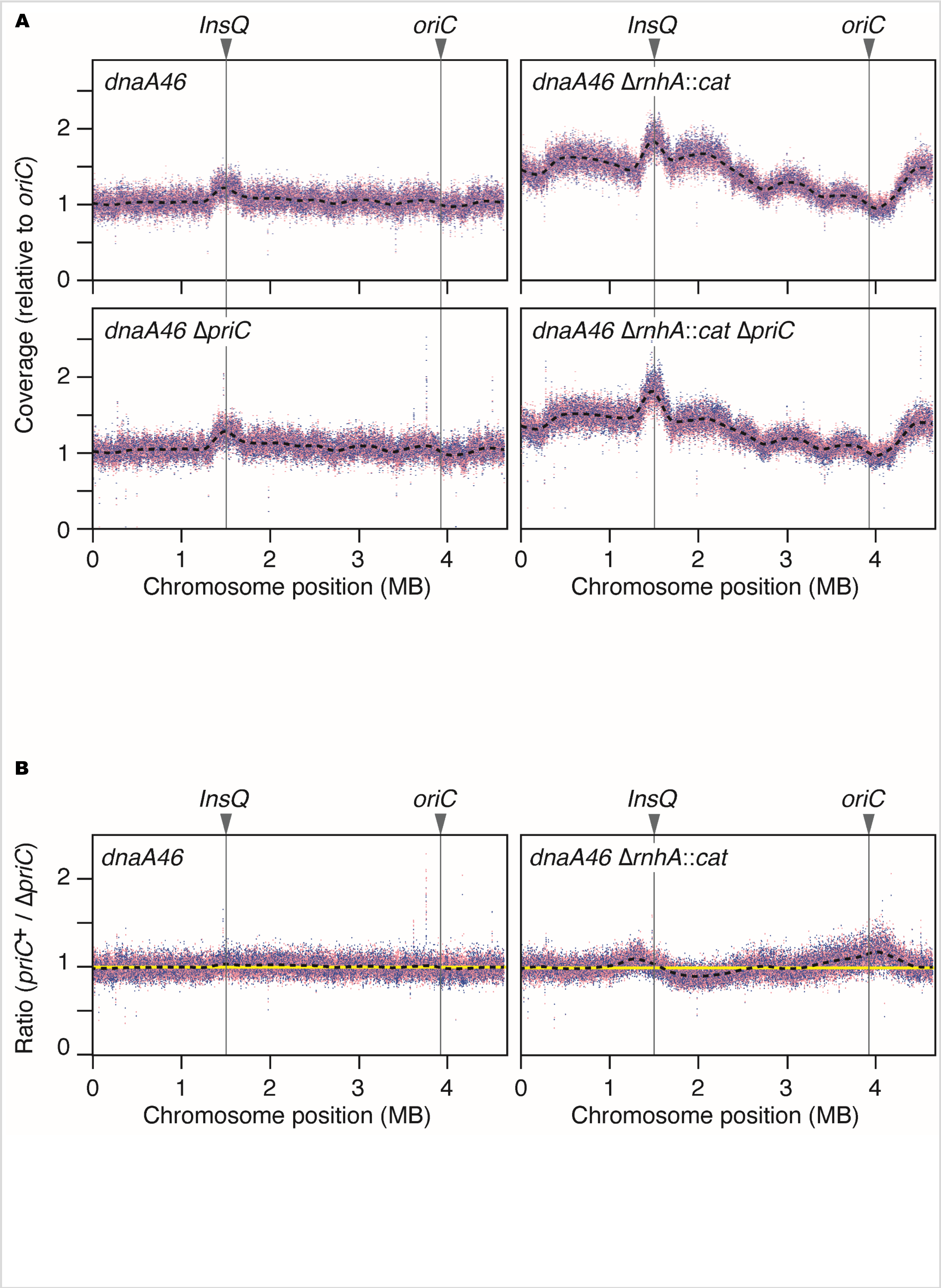
